# Quantifying posterior effect size distribution of susceptibility loci by common summary statistics

**DOI:** 10.1101/714287

**Authors:** Olga A. Vsevolozhskaya, Dmitri V. Zaykin

## Abstract

Testing millions of SNPs in genetic association studies has become standard routine for disease gene discovery, followed by prioritization of the strongest signals based on the set of the smallest P-values. In light of recent re-evaluation of statistical practice, it has been suggested that P-values are unfit as summaries of statistical evidence. Despite this criticism, P-values are commonly used and are unlikely to be abandoned by practitioners. Moreover, P-values contain information that can be utilized to address the concerns about their flaws and misuse. We present a new method for utilizing evidence summarized by P-values for estimating odds ratio (OR) based on its approximate posterior distribution. In our method, only P-value, sample size, and standard deviation for log(OR) are needed as summaries of data, accompanied by a suitable prior distribution for log(OR) that can assume any shape. The parameter of interest, log(OR), is the only parameter with a specified prior distribution, hence our model is a mix of classical and Bayesian approaches. We show that our “Mix Bayes” (MB) method retains the main advantages of the Bayesian approach: it yields direct probability statements about hypotheses for OR and is resistant to biases caused by selection of top-scoring SNPs. MB enjoys greater flexibility than similarly inspired methods in the assumed distribution for the summary statistic and in the form of the prior for the parameter of interest. We illustrate our method by presenting interval estimates of effect size for reported genetic associations with lung cancer. Although we focus on OR, our method is not limited to this particular measure of effect size and can be used broadly for assessing reliability of findings in studies testing multiple predictors.

## Introduction

Modern human genetics studies routinely examine very large numbers of potential associations of genetic variants with health related outcomes. In the majority of studies, results are filtered by P-values with the most promising results selected based on a significance threshold, adjusted to accommodate the number of tests in a study. Partly due to their widespread use, P-values have been at the center of replicability crisis. ^1–3^ P-values have high study-to-study variability and they fail to measure confidence in a hypothesis, even though the operational use of P-values is to make decisions about hypotheses. ^4–7^ Although P-values are poorly suited for what they are used for in practice, they are efficient summaries of data, and they contain information that can be used to judge degree of uncertainty about hypotheses and parameters. The Bayesian framework gives researches the ability to address trustworthiness of their hypothesis directly. There are a number successful proposals where information summarized by P-values is used for Bayesian or approximately Bayesian inference. The false discovery rate (FDR) is perhaps the most well known example. The FDR is defined as an estimated proportion of false discoveries among test results that are deemed to be “discoveries,” based on statistical criteria. The FDR has distinct Bayesian characteristics because the posterior probability of a hypothesis, averaged over discoveries, is the proportion of false discoveries estimated for a given study. The original FDR method by Benjamini and Hochberg is not a truly Bayesian approach, nor was it intended to be one by design, because the FDR in this method is controlled as a handily conceived long run average, taken over experiments with or without discoveries. When statistical power is high (and hence assuming that not all of study’s null hypotheses are true), Benjamini-Hochberg’s FDR can be viewed as a proportion of false findings estimated concerning *a specific study*. That is, conveniently, as sample size increases, Benjamini-Hochberg’s FDR approaches the Bayesian posterior proportion of true null hypotheses among rejected hypotheses. Likewise, based solely on P-values are Empirical Bayes counterparts of the Benjamini-Hochberg FDR, such as Storey’s q-value^8^ and the local FDR.^9^ These two methods preserve the ranking of P-values, for example, the order of q-values is the same as that of P-values, and thus, q-values can be viewed as posterior summaries concerning the standardized effect size (e.g., estimated logarithm of odds ratio divided by its estimated standard deviation).

The Empirical Bayes methods became popular in applications such as the analysis of differential gene expression. However, the applicability of Empirical Bayes approaches for genome-wide association studies is limited. The reason behind this limitation is the fact that Empirical Bayes methods use data at hand to infer prior parameters, but genome-wide studies tend to have relatively weak signals, which makes it challenging to reliably estimate prior parameters from the same data that will be used for subsequent Bayesian analysis.^10^ Nonetheless, current knowledge of genetics of complex diseases may allow one to specify plausible prior parameters explicitly. For example, it would be reasonable to assume that there is a very small *a priori* chance that a randomly picked SNP would carry OR value greater than 3, and that probability of OR value being at least 2 is also small, but definitely larger. Further, one can place with high confidence a large prior probability for the OR to be around 1. Thus, a biologically realistic prior distribution for an effect size can be constructed based on existing prior knowledge.

A simple method that accommodates explicit prior assumptions is the False Positive Report Probability (FPRP) by Wacholder et al.^11^ The method received great attention and has been cited over 1500 times at the time of this writing, in part due to its simplicity. In the FPRP, the prior proportion of non-associated SNPs, Pr(*H*_0_), should be specified, as well as a prior value for OR. That is, a single OR value is used to approximate the actual OR distribution across associated variants. Given these two assumed values, power can be estimated for a given SNP. Then, FPRP/(1-FPRP) can be calculated as the ratio of P-value over the computed power times the prior odds for *H*_0_. The FPRP method has a number of shortcomings. Among the main ones is the assumption that a single value of OR is representative of the OR distribution, and the use of cumulative probabilities (P-value and power), which leads to an undesirable property that the FPRP can never be greater than the assumed prior probability of *H*_0_.^12,13^

Wakefield^14^ proposed a Bayesian method that is similar in simplicity to FPRP but remedies its shortcomings. Wakefield’s Bayesian measure of the probability of false discovery, called the approximate Bayes factor (ABF), utilizes information contained in a test statistic for ln(OR) to obtain an approximate posterior effect size distribution for a given SNP. To approximate the posterior effect size distribution for real signals, the ABF method assumes that the effect size distribution across the genome, measured by the logarithm of odds ratio, can be approximated by a normal distribution centered at zero. Large values of the variance parameter, *W*, of that prior distribution imply large spread of ln(OR) values, and therefore, high probability of encountering variants with relatively large effect size. Non-associated variants in ABF are modeled as having zero effect size. Alternatively, if for non-associated SNPs we assume a normal distribution with nearly zero variance, then the prior for both real and non-associated variants becomes a mixture of two normal distributions, both centered at zero. Thus, the prior distribution in ABF is controlled by two parameters, the mixture proportion of non-associated SNPs, Pr(*H*_0_), and the variance of the effect size distribution for associated SNPs, *W*.

The ABF method and its extensions^15^ are “approximate Bayes” methods, because the posterior calculation, which is based on a *Z*-statistic for ln(OR), assumes the prior distribution for the parameter of interest only, i.e., ln(OR). In contrast, a fully Bayesian model would have to include a joint prior distribution for covariates of the model along with ln(OR). It can be difficult to specify a realistic joint prior for all of the parameters, and the ABF finds a middle ground in which approximate, yet surprisingly precise posterior estimates for ln(OR) are obtained by utilizing its prior distribution and by plugging in frequentist estimates for any additional parameters. The ABF has desirable statistical properties, including explicit drawing on power through dependence on the standard error. Note that P-values are calculated assuming *H*_0_, and under this assumption they are not a function of the sample size. This property of ABF leads to a generally different ranking than ranking by P-values and facilitates discovery of real signals.^13^

A great practical advantage of the ABF is that only ln(OR), the corresponding normal *Z*-statistic, and the standard error of ln(OR) are needed as summaries of data. In this paper, we propose a method that similarly requires only these summaries of data but is not limited to OR as a measure of effect size and offers much greater flexibility in terms of the prior distribution for ln(OR). Similarly to ABF, our “Mix Bayes” approach improves precision of effect size estimates for multiple SNPs tested within a study and makes use of summary statistics to estimate the posterior distribution for the parameter of interest. However, the core novelty of our approach is the ability for a researcher to use any arbitrary prior effect size distribution, e.g., based on previously published empirical results. Our method includes ABF as a special case when the conjugate normal model is assumed, therefore it shares ABF’s advantageous statistical properties demonstrated previously by Wakefield.^13,14,16^ The ability to specify an arbitrary prior distribution in our method is important, because a well-specified (prior) distribution for effect sizes in the genome makes the posterior effect size estimates resilient to bias due to selection of top-ranking results. For example, one can take a SNP with the smallest P-value in a GWAS and compute the posterior probability of *H*_0_. This posterior probability, as well as estimates of effect size, is unaffected by multiple testing or by selection of top ranking results, provided the prior effect size distribution is specified realistically. In contrast, the usual estimates, e.g., the largest odds ratios observed in a study, tend to be over-estimating the actual values – a phenomenon known as the winner’s curse. Comparing to P-value adjustments in multiple testing, this advantage of the Bayesian approach is more than a simple convenience, because it is not always straightforward to define the amount of testing that a P-value needs to be adjusted for, and even a properly adjusted P-value still lacks probabilistic interpretation in terms of degree of confidence that the result is not spurious. ^1^

Robustness of posterior estimates to selection bias holds only when the effect size distribution is specified correctly.^17^ One concern is that the normal prior distribution assumed in ABF may not always provide sufficient flexibility. For example, one may prefer an asymmetric prior distribution because without randomization of which allele is being tested at each SNP, the symmetry implies that susceptibility and protective effects are equally likely. Further, expecting that the bulk of truly associated SNPs in genome-wide studies carry small effect sizes, it is desirable to be able to accommodate distributions where there are occasional moderate or large effect sizes, while a sizable part of the density is closer to zero than what a normal distribution can provide. Moreover, methods have been emerging for estimation of disease-specific effect size distributions from GWAS and replication studies. These methods allow one to utilize effect size distributions reported in a tabulated way, where each range of the effect size would be accompanied by its estimated frequency in the genome. Such empirically estimated distributions are not necessarily expected to follow any standard or symmetric form.

Via application of the Mix Bayes approach to simulated and real data, we found that Mix Bayes produces unbiased effect size estimates under various forms of selection. Selection is not only expected to generate bias in classical effect estimates but this bias is difficult to correct for. While many adjustments for the winner’s curse bias have been proposed, there can be no unbiased estimate for the top-ranking hits^18^ unless external information, which takes the form of a prior distribution in Bayesian approaches, is utilized. Our method incorporates this prior information in a flexible way and shares simplicity of ABF and FPRP, requiring only summary statistics for its implementation.

## Methods

Consider test statistics that can be expressed as a product of the sample size and the standardized effect size, i.e., effect size scaled by variance. These statistics span the majority of tests used for genetic association analyses, including normal, chi-squared, Student’s *t*, and F densities. For instance, a standard *Z*-score, 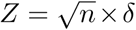, satisfies this property, where *δ* = *µ/σ* is the standardized effect size. If the outcome is a case/control classification with a binary predictor (exposure), the test statistic can be expressed as:

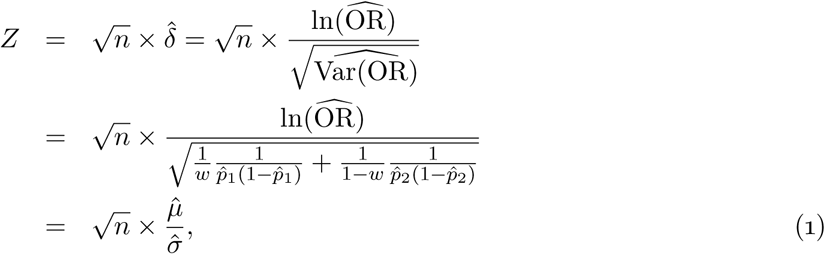

where *w* is the proportion of cases in the sample, 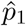 is the frequency of exposure among cases and 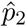 is that among the controls. For these statistics, deviations from the null hypothesis can be expressed by a single parameter *γ*. For example, under the alternative hypothesis, the standard score and the statistic in Eq. (1) will be distributed as *Z* ∼ *N* (*γ*, 1), where 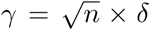 is the non-centrality parameter.

In our method, posterior distribution for the effect size attributed to a genetic variant is first obtained in the standardized form, from which approximate posterior distribution for ln(OR) can be obtained as follows. We start by supplying some prior (external) information regarding possible values of the effect size for *µ*=ln(OR), with their respective probabilities. For example, one may consider testing whether a SNP has an effect size that is at most |*µ*^0^| in magnitude and define the null hypothesis *H*_0_ : *µ ≤* |*µ*^0^| with the alternative hypothesis specified as *H*_*A*_ : *µ >* |*µ*^0^|. Then, the proportion of SNPs with effect sizes smaller than |*µ*^0^| can be regarded as the prior probability of *H*_0_, *π* = Pr(*H*_0_), and Pr(*H*_*A*_) = 1 *-π*. This specification makes a sharp distinction between effect sizes that are large enough to be considered genuine and correspond to *H*_*A*_, and a set of smaller effect sizes that correspond to *H*_0_, and can be represented by a mixture of two distributions with the mixture proportion *π*. A more flexible way is to model all tested signals together as arising from an effect size distribution, without specifying the form of *H*_0_ upfront. For squared values of effect sizes, e.g., ln^2^(OR), such distribution may be an L-shaped, finely binned histogram with a sizable spike around zero that reflects a preponderance of signals carrying small effects. In terms of genome wide association scans, such distribution reflects prior knowledge that a randomly chosen SNP has a small effect size with high probability. Under this model, instead of specifying priors for the dichotomous scenario as *π* and 1 *-π*, we would partition the prior distribution of *µ* into a finite set of values *µ*_1_, *µ*_2_, …, *µ*_*B*_ with the corresponding prior probabilities, Pr(*µ*_*i*_). Then, noting one-to-one correspondence between P-value and Z-statistic, the posterior probability for the standardized effect size to take the value *δ*_*j*_ is

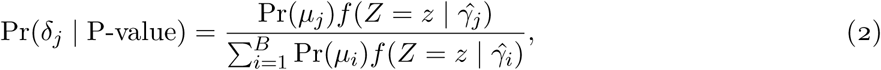

where *f* (·) is the test statistic density with the non-centrality parameter 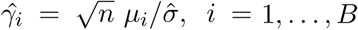, and *z* is the observed value of the test statistic *Z*. Once all posterior probabilities corresponding to each value of *γ* are obtained, the values 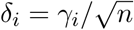 form a finely binned histogram of the posterior distribution for the standardized effect size. With this posterior distribution, one can construct (1−*α*)% credible intervals, make probabilistic statements, and estimate the standardized effect size as the posterior mean:

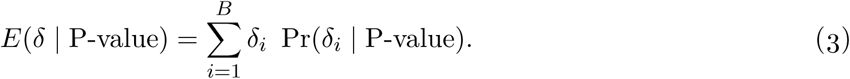

Utility of standardized effect sizes or standardized regression coefficients (*δ*) may be questioned,^19–22^ however, posterior distribution for the standardized parameter can be converted to an approximate posterior distribution of the parameter itself (*γ*), by plugging in the sample standard error, e.g., 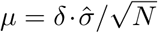. With our focus on odds ratio, disease risk *p* is related to exposure and other variables via logistic model, logit(*p*) = **x**^′^***β*** + *zµ*, where **x**^′^ and ***β*** are vectors of covariates and their respective coefficients, and *µ* is ln(OR) with the corresponding exposure value, *z*. Following Wakefield,^14^ we consider a prior distribution for *µ* only, rather than a joint prior on all parameters.

## Results

### Simulation study and methods comparisons

We first assessed the performance of the proposed method through a simulation study, with the following objectives: (1) to show that the approximate posterior distribution for ln(OR) can be accurately extracted from P-value, and (2) to show that the resulting posterior distributions is robust to selection bias in large scale experiments. The Mix Bayes approach allows one to obtain effectively exact posterior distribution for the standardized effect size and an approximated posterior distribution for the effect size itself.

In a special case of the conjugate normal model with zero-centered normal prior, MB is expected to approach ABF as the number of bins of the discretized normal prior increases. Although in this case we expect posterior estimates obtained by two methods to be the same, it is of interest to confirm their resistance to the winner’s curse. Further, it is critical for a posterior approximation to be robust to various forms of selection. For example, we want to make sure that the effect size is not going to be overestimated if a predictor with the maximum observed OR or the minimum P-value is selected from a multiple testing experiment. When prior information is specified correctly, posterior estimates should be unaffected by bias due to selection. The key advantage of the method compared to P-values is that posterior estimates (e.g., of odds ratios) and probabilities of hypotheses do not need adjustments for multiple testing.

To generate data under the normal prior model (where ABF and MB are expected to be equivalent), we assumed that among 20,000 SNPs there would be at most one with an odds ratio over two, and from that we determined the prior variance *W* = 0.03174. Thus, the prior distribution for the effect size was ln(OR) ∼ *N* (0, *W*). For the implementation of the Mix Bayes approach purposes, we discretized this distribution to 3,387 bins (this value is obtained by truncating the prior normal distribution at its 10^*-*6^ and 1 *-* 10^*-*6^ quantiles and using the bins of length 5 × 10^*-*4^. Further details of data generation step are given in Appendix (Simulation Study Setup section).

For each simulation, we performed a total of *L* tests and selected top-ranking SNPs based on one of the “selection rules”, such as P-value threshold. After calculating the estimated 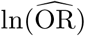 and the posterior expectations for the selected SNP, we averaged the results across simulations and obtained (1) the expected values of the true log odds ratio that gave rise to the selected test statistic, *E*(ln(OR)), (2) the corresponding average of posterior expectations, and (3) the averaged frequentist estimate for the log of odds ratio,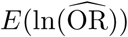.

Table 1 presents the results of SNP selection based on either the minimum P-value or the maximum ln(OR) absolute value. These types of selection are not only expected to generate bias in classical effect estimates (the winner’s curse phenomenon) but are also difficult to correct for. While many adjustments for the winner’s curse were previously proposed, there is no unbiased estimate for the top-ranking hits in the classical (frequentist) framework without the usage of external information, which takes the form of a prior distribution in Bayesian approaches.^18^ As Table 1 clearly indicates, ABF and Mix Bayes posterior estimates are almost identical and are unbiased with respect to the true expected value *E*(ln(OR)), computed as the average of the actual values of ln(OR) across simulations. This average is not equal to zero (the mean of the prior distribution) due to selection of top-ranking results, as reflected by the first column of Table 1. Further, the last column of Table 1 indicates that the simple average over the maximum of frequentist estimates of ln(ORs), 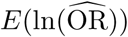, is highly biased for small sample sizes (*n* = 500) and moderately biased for larger sample sample sizes (*n* = 5000). Thus, relative to the classical approach, Bayesian approaches require substantially smaller sample sizes to produce unbiased estimates of the effect size of the top-ranked SNPs.

**Table 1:** Summary of ln(OR) expectations out of *L* tests.

| # of tests ($L$) | n | True value | Posterior estimate | | Frequentist estimate |
| --- | --- | --- | --- | --- | --- |
| | | $E(\ln(\text{OR}))$ | ABF | Mix Bayes | $E(\ln(\widehat{\text{OR}}))$ |
| Select min P |  |  |  |  |  |
| 1,000 | 500 | 0.38 | 0.38 | 0.38 | 0.85 |
|  | 1,000 | 0.45 | 0.45 | 0.44 | 0.71 |
|  | 5,000 | 0.53 | 0.53 | 0.53 | 0.59 |
| $1 \times 10^5$ | 500 | 0.53 | 0.51 | 0.51 | 1.12 |
|  | 1,000 | 0.61 | 0.60 | 0.60 | 0.95 |
|  | 5,000 | 0.72 | 0.72 | 0.71 | 0.90 |
| Select max OR |  |  |  |  |  |
| 1,000 | 500 | 0.33 | 0.31 | 0.31 | 0.95 |
|  | 1,000 | 0.42 | 0.41 | 0.41 | 0.77 |
|  | 5,000 | 0.53 | 0.54 | 0.54 | 0.62 |
| $1 \times 10^5$ | 500 | 0.37 | 0.32 | 0.32 | 1.40 |
|  | 1,000 | 0.51 | 0.48 | 0.48 | 1.09 |
|  | 5,000 | 0.73 | 0.73 | 0.71 | 0.84 |

A different type of selection is when SNPs are selected based on a P-value magnitude. For example, “P ∼ 0.05” in the first column of the Table 2 denotes the type of selection where P-values that fall in a narrow interval around 0.05 were retained (0.05±0.05/2). The second part of the table shows a thresholding type of selection, that is, selection of SNPs with P-values that are smaller than a given threshold. On the one hand, average of frequentist estimates of ln(OR) across simulations, 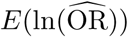, shows bias due to selection – a consequence of the winner’s curse. On the other hand, ABF and Mix Bayes approaches produce unbiased values that are expectedly very similar to one another. No correction for multiple testing was done to either the ABF or the Mix Bayes method, nor it is needed, because the prior distribution already properly accommodates the high frequency of SNPs carrying very small values of ln(OR).

**Table 2:** Expectations of ln(OR) under different types of P-value selection.

| Type of selection | n | True value | Posterior estimate |  | Frequentist estimate |
| --- | --- | --- | --- | --- | --- |
| | | $E(\ln(\text{OR}))$ | ABF | Mix Bayes | $E(\ln(\widehat{\text{OR}}))$ |
| $P \sim 0.05$ | 1,000 | 0.17 | 0.17 | 0.17 | 0.29 |
| $P \sim 0.001$ | 1,000 | 0.29 | 0.29 | 0.29 | 0.47 |
| $P \sim 1 \times 10^{-5}$ | 1,000 | 0.38 | 0.38 | 0.38 | 0.62 |
| $P \sim 1 \times 10^{-7}$ | 1,000 | 0.46 | 0.46 | 0.46 | 0.73 |
| $P < 0.05$ | 1,000 | 0.24 | 0.24 | 0.24 | 0.39 |
| $P < 0.001$ | 1,000 | 0.34 | 0.33 | 0.33 | 0.54 |
| $P < 1 \times 10^{-5}$ | 1,000 | 0.42 | 0.42 | 0.42 | 0.68 |
| $P < 1 \times 10^{-7}$ | 1,000 | 0.50 | 0.49 | 0.49 | 0.78 |

Based on posterior distribution of ln(OR), one can also obtain credible interval estimates, as provided in Table 3. It is interesting to compare posterior intervals to the classical confidence intervals (CI). By their construction, CI’s incorporate the implicit assumption that the variance of the prior distribution is so large that any value of ln(OR) is equally likely. The influence of this assumption on the interval endpoints decreases with the sample size, *n*, and for that reason it may be incorrectly assumed that genetic studies with thousands of observations would yield intervals which endpoints can be interpreted as probabilistic bounds for ln(OR). However, due to sparsity of SNPs carrying large effect sizes as well as to selection of top-ranking results, CIs can remain substantially biased even with large *n*. In contrast, coverage of the ABF and the Mix Bayes methods is around the declared 80%. The type of selection shown in the table is P-value-based. Similar results were obtained for selections based on large values of odds ratios instead of P-values (results not shown).

**Table 3:** Empirical coverage of the two credible intervals and a confidence interval at the 80% nominal level

| Type of Selection | $n$ | 80% Empirical Coverage | | |
| --- | --- | --- | --- | --- |
|  |  | ABF | Mix Bayes | CI |
| $P \sim 0.05$ | 1,000 | 79.90% | 79.82% | 72.70% |
| $P \sim 0.001$ | 1,000 | 80.39% | 80.33% | 50.13% |
| $P \sim 1 \times 10^{-7}$ | 1,000 | 80.29% | 79.94% | 20.60% |
| $P < 0.05$ | 1,000 | 79.85% | 79.78% | 60.12% |
| $P < 0.001$ | 1,000 | 80.22% | 80.11% | 41.13% |
| $P < 1 \times 10^{-7}$ | 1,000 | 80.37% | 79.40% | 16.56% |
| min P, 1K tests | 500 | 80.43% | 80.12% | 7.91% |
|  | 5,000 | 79.82% | 78.99% | 60.43% |
| min P, 10K tests | 500 | 80.27% | 79.26% | 2.38% |
|  | 5,000 | 80.76% | 79.98% | 55.28% |

One of the features of the proposed Mix Bayes approach is its flexibility with regard to what shape the prior distribution can take. To illustrate this feature, we used an asymmetric prior distribution shown in Figure 1 to obtain simulation results for top SNPs selected based on the minimum P-value in experiments with *L* =1,000, 50,000, and 10^5^ tests. The results are summarized in Table 4 and show that posterior expectations calculated with the Mix Bayes approach closely match the true average values of the effect size, while estimated values from the logistic regression overestimate the effect size due to the winner’s curse. Finally, we checked the spread of Mix Bayes posterior expectations relative to the true value of ln(OR), given the asymmetric prior. Figure 2 shows contour plots of the Mix Bayes posterior estimates for ln(OR) against the actual values of ln(OR). The classical estimates, 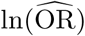 from the equation for the *Z*-statistic were also included. Each point on the graphs was selected based a SNP with the largest ln(OR) out of 10,000 tests. As expected, compared to the Mix Bayes method, the classical estimate exhibit the winner’s curse, i.e., upward bias, as well as a lower correlation with the true value of 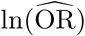 (i.e., plots for the classical estimates appear higher on the graphs and they are less elliptical than those for the Mix Bayes estimates). The two types of estimates become more similar as sample size increases.

**Figure 1:**
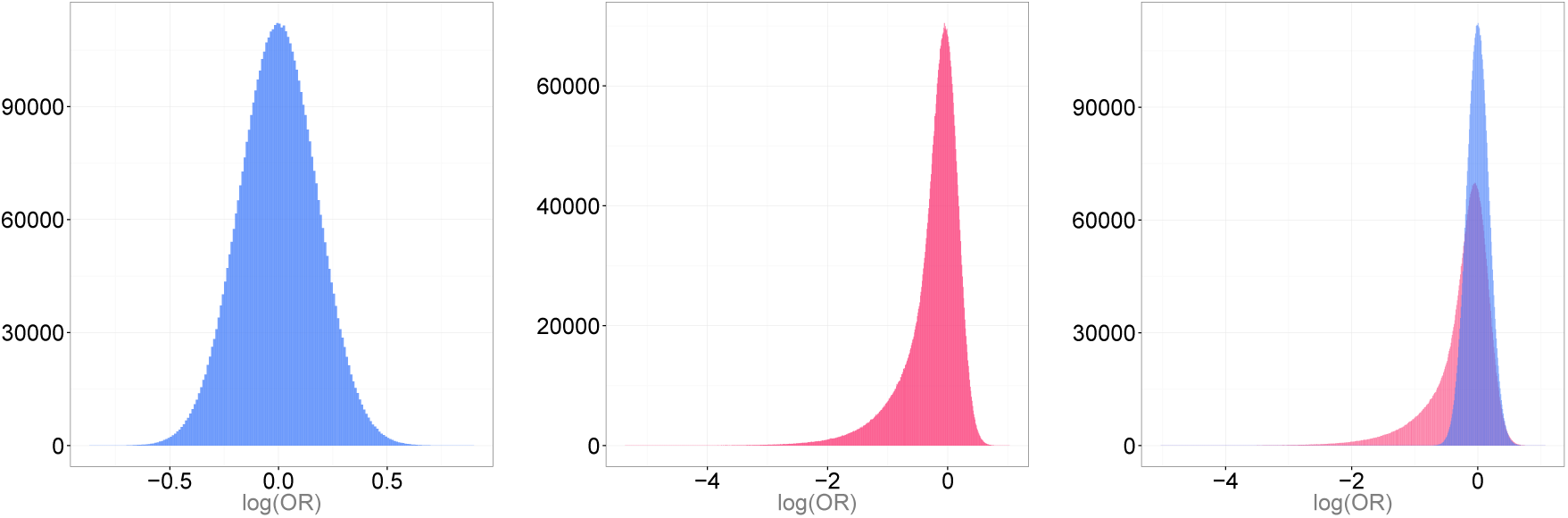
Conjugate Normal and Non-conjugate Skewed Prior Distributions for log(OR) The left graph is a density of the normal prior with the variance set to approximate the occurrence of OR>2 at about 1 in 20,000 SNPs. The middle graph is visibly asymmetric and represents non-conjugate skewed prior. The right graph plots the two distributions on the same scale with the overlap shown in purple.

**Figure 2:**
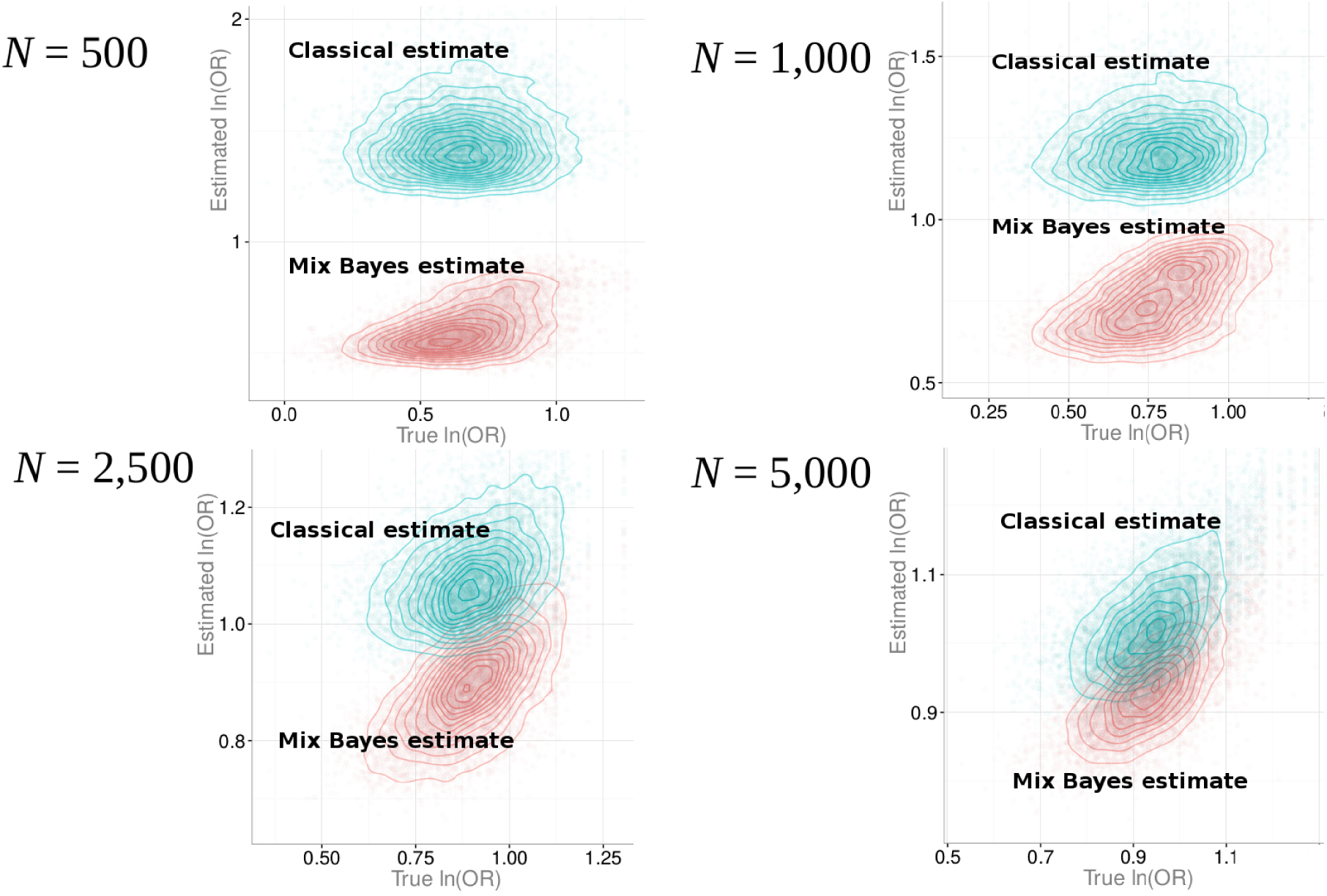
Simulation scatter plots. Each point in the graphs is an association result selected on the basis of a SNP with the minimum P-value out of 10,000 tests.

**Table 4:** Summary of ln(OR) expectations for the minimum P-value out of *L* tests with a non-conjugate prior distribution

| # of tests ($L$) | $n$ | $E(\ln(\text{OR}))$ | Mix Bayes | $E(\ln(\widehat{\text{OR}}))$ |
| --- | --- | --- | --- | --- |
| 1,000 | 500 | 0.67 | 0.67 | 1.08 |
|  | 1,000 | 0.74 | 0.73 | 0.96 |
|  | 5,000 | 0.81 | 0.81 | 0.86 |
| 50,000 | 500 | 0.83 | 0.81 | 1.35 |
|  | 1,000 | 0.90 | 0.90 | 1.20 |
|  | 5,000 | 1.01 | 1.01 | 1.07 |
| $1 \times 10^5$ | 500 | 0.85 | 0.84 | 1.39 |
|  | 1,000 | 0.93 | 0.92 | 1.24 |
|  | 5,000 | 1.03 | 1.03 | 1.10 |

**Table 5:** Reanalysis of variants associated with lung cancer risk.

| Chr | Gene | Variant ID | Major/Minor<br>Allele | Stage | Case | Con | MAF |  | OR (95% CI) | P-value |
| --- | --- | --- | --- | --- | --- | --- | --- | --- | --- | --- |
|  |  |  |  |  |  |  | Case | Con |  |  |
| 6p21.33 | *BAT2* | rs9469031 | C/T | Disc | 1,341<sup>a</sup> | 1,962<sup>a</sup> | 0.019 | 0.036 | 0.52 (0.37-0.73) | 1.54<sup>a</sup> × 10<sup>-4</sup> |
|  |  |  |  | Rep I | 1,114 | 1,245 | 0.024 | 0.042 | 0.61 (0.44-0.83) | – |
|  |  |  |  | Rep II | 3,508 | 3,631 | 0.011 | 0.018 | 0.62 (0.46-0.84) | – |
|  |  |  |  | Post (mean) | – | – | – | – | 0.68 (0.43-1.07) | – |
|  |  |  |  | Post (med) | – | – | – | – | 0.62 (0.43-1.01) | – |
| 6p21.33 | *FKBPL* | rs200847762 | G/A | Disc | 1,341<sup>a</sup> | 1,982<sup>a</sup> | 0.004 | 0.19 | 0.21 (0.11-0.39) | 1.84<sup>a</sup> × 10<sup>-6</sup> |
|  |  |  |  | Rep I | 1,110 | 1,239 | 0.003 | 0.014 | 0.19 (0.08-0.46) | – |
|  |  |  |  | Rep II | 3,581 | 3,666 | 0.002 | 0.003 | 0.66 (0.34-1.28) | – |
|  |  |  |  | Post (mean) | – | – | – | – | 0.27 (0.13-0.55) | – |
|  |  |  |  | Post (med) | – | – | – | – | 0.27 (0.13-0.54) | – |
| 6p22.2 | *HIST1H1E* | rs2298090 | A/G | Disc | 1,341<sup>a</sup> | 1,982<sup>a</sup> | 0.006 | 0.020 | 0.32 (0.19-0.56) | 6.16<sup>a</sup> × 10<sup>-5</sup> |
|  |  |  |  | Rep I | 1,101 | 1,235 | 0.013 | 0.024 | 0.56 (0.36-0.87) | – |
|  |  |  |  | Rep II | 3,429 | 3,551 | 0.006 | 0.008 | 0.67 (0.44-1.02) | – |
|  |  |  |  | Post (mean) | – | – | – | – | 0.48 (0.24-1.01) | – |
|  |  |  |  | Post (med) | – | – | – | – | 0.42 (0.18-1.00) | – |
| 20q11.21 | *BPIFB1* | rs6141383 | G/A | Disc | 1,341<sup>a</sup> | 1,982<sup>a</sup> | 0.025 | 0.012 | 2.00 (1.36-2.95) | 4.80<sup>a</sup> × 10<sup>-4</sup> |
|  |  |  |  | Rep I | 1,110 | 1,240 | 0.022 | 0.013 | 1.68 (1.07-2.63) | – |
|  |  |  |  | Rep II | 3,507 | 3,455 | 0.019 | 0.013 | 1.64 (1.24-2.17) | – |
|  |  |  |  | Post (mean) | – | – | – | – | 1.31 (0.73-2.36) | – |
|  |  |  |  | Post (med) | – | – | – | – | 1.01 (0.43-2.36) | – |

Abbreviations are as follows. Chr: chromosomal region; MAF: minor allele frequency; CI: confidence interval; Disc: discovery; Rep: replication; Post: posterior; Con: Control; med: median.
<sup>a</sup> Summary values from the discovery study that were used to calculate posterior expectations and 95% credible intervals.

In summary, our results reassure the validity of the proposed approximate Bayesian approach, where the prior distribution is specified only for the parameter of interest. For example, a potential concern could have been that the odds ratio alone is not sufficient to describe a population in terms of parameters that affect disease risk. Risk of disease given allele, Pr(*D*|*A*), and allele frequency, Pr(*A*), are two other parameters that also have respective distributions across SNPs. Thus, “simulated reality” in our experiments was derived from three random distributions rather than from a single one for ln(OR). In contrast, during the analysis of simulated data by our method, a prior distribution is specified for the ln(OR) only. The three random parameters (ln(OR), Pr(*D*|*A*), and Pr(*A*)) together determine prevalence of disease due to SNP as well as allele frequencies among affected and unaffected individuals. Furthermore, standard deviation for ln(OR) is also random, as a function of these parameters. Therefore, Pr(*D*|*A*) and Pr(*A*) would have to be part of a prior distribution in a fully Bayesian model. Nevertheless, our simulation studies confirm validity of the proposed method as judged by unbiased point and interval posterior estimates in the presence of potentially bias-inducing selection of top-scoring P-values or odd ratios.

### Lung cancer association study

We applied the Mix Bayes approach in re-analysis of four rare candidate variants reported by Jin et al.^23^ in a study of low-frequency genetic associations with lung cancer. To calculate posterior effect sizes for candidate susceptibility variants, we used their reported P-values from the genome-wide discovery stage^23^ and prior effect size distribution for cancers, as reported by Park et al.^24^ Genetic variant with the smallest discovery P-value reported by Jin et al.^23^ was rs200847762 within the *FKBPL* gene (OR = 0.21, P = 1.84×10^*-*6^). This variant replicated in first of the two replication studies (OR = 0.19, 95% Confidence Interval: (0.08 – 0.46)). In the second replication study, the estimated OR was consistent with the discovered effect direction, 0.66, but the confidence interval included one (0.34 – 1.28). Application of the Mix Bayes approach produced the posterior estimate of OR for the discovery study equal to 0.27, with the 95% Credible Interval (0.133-0.55), suggesting a genuine causal signal within this region (we also report posterior estimates for the median of OR, which are close in values to the estimates of the posterior mean). This signal was further suggested to be in association with breast cancer among Chinese women by Zhou et al.^25^ The therapeutic and diagnostic potential of *FKBPL* to targeting tumours was discussed by Robson and James^26^ and McKeen et al.^27^ Genetic variant with the second smallest P-value in Jin et al. discovery study was rs2298090 within the *HIST1H1E* gene (P = 6.16×10^*-*5^). This signal also failed to replicate at the second stage of their study, OR = 0.65, 95% CI (0.44 – 1.02). The Mix Bayes posterior mean estimate was OR = 0.48 with the 95% interval (0.24-1.01). Finally, rs9469031 within the *BAT2* gene (also known as *PRRC1A*) and rs6141383 within the *BPIFB1* gene (both discovery P-values ≈ 10^*-*4^) were replicated in the second stage of the original study by Jin et al.^23^ The Mix Bayes estimates suggest lower effect size magnitudes than what is reported in the discovery stage. Specifically, for rs9469301 candidate SNP, OR = 0.68, with 95% posterior interval (0.43 – 1.07); for rs6141383, OR = 1.31, with 95% posterior interval (0.73 – 2.36). The posterior OR estimates tend to be closer to the replication values than to the possibly inflated values found during the discovery stage. Note that in all cases posterior intervals are wider than the 95% confidence intervals for the discovery stage, because Bayesian intervals account for sparsity of real associations via the prior distribution specified for OR and thus do not need adjustment for multiple testing.

## Discussion

Ongoing “replicability crisis” put P-values at the center of controversy. Although P-values are often misused and misinterpreted, they are routinely available and reported. In this article, we suggest a new method to compute an approximate posterior distribution for ln(OR), in part based on P-value or the usual test statistic for testing significance of odds ratio. This posterior distribution can be used in versatile ways. One can estimate posterior probability of genetic association with disease, obtain point estimates and test interval hypotheses about probabilistic ranges for the effect size. The method is unaffected by the winner’s curse and by multiple testing, provided one can realistically describe the effect size distribution across SNPs in the genome.

Our method is “approximate” in reference to its Bayesian part, because the prior distribution is specified only for effect size, the parameter of interest. Wakefield’s ABF^14^ is approximate in the same sense as our method. In other words, approximation is a compromise between Bayesian and classical approaches. In a fully Bayesian model, complete joint prior distribution for all parameters (including coefficients for covariates) should be specified, leading to increased computational complexity, and to difficulties in specification of prior distributions with realistic dependencies between parameters. The approximate Bayesian methods enable one to utilize our basic understanding about the distribution of effect size (such as relative risk or odds ratio) across genetic variants in the genome and to construct posterior estimates from simple and readily available summary statistics. Through simulation studies, we confirmed effectiveness of this “compromise” strategy, where we place a prior distribution on the effect size only and demonstrated that the proposed method is resistant to potential bias due to selection of top-ranking P-values and odds ratios and that it gives accurate point and interval estimates of true effect size.

The proposed Mix Bayes approach is general in the sense that it can be used with any prior effect size distribution and with statistics other than the normal Z-score for ln(OR). One illustration is given in the link to our software below, where we implemented a simulation example for testing validity of posterior inference for the mean difference between two groups, using the two-sample T-test statistic as a summary of data. Parameters of the prior distribution in our method, e.g., prior variance for the distribution of log(OR), do not need to be specified directly, because the prior distribution can be handcrafted as a table of values with their respective frequencies. This can be done with relative ease as methods emerge for estimating effect size distribution from genome-wide association data.^24,28^ Such empirically estimated distributions are not necessarily expected to follow any symmetric or standard form. Thus, more flexibility is needed, and the Mix Bayes method allows researchers to incorporate empirically estimated distributions directly, without having to choose a prior from one of the standard distributions. That being said, any standard distribution can be easily discretized and used as a prior in applications of our method. Another obvious application is re-evaluation of published findings. Access to individual records may be difficult or impossible, but summary statistics needed for approximate posterior inference are usually available. Posterior distributions for standardized effect sizes are particularly simple and expected to be accurate when such distribution is derived from a test statistic whose non-centrality is a function of standardized effect size.

## Web Resources

The URL for software referenced in this article is available at:

https://github.com/dmitri-zaykin/Mix_Bayes

## Acknowledgements

This research was supported in part by the Intramural Research Program of the NIH, National Institute of Environmental Health Sciences.

## Appendix

### ABF: A special case of Mix Bayes

Although our proposed Mix Bayes approach is very general in the sense that it can deal with any arbitrary prior effect size distribution, additional insights into its behaviour may be gained by showing its equivalence to the approximate Bayes factor^14^ under the conjugate normal model. Specifically, Wakefield^14^ showed that if disease risk, *p*, is modeled via logistic regression, then assuming a zero-centered normal prior distribution for *µ* = ln(OR) ∼ *N* (0, *W*) and given 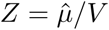, where *V* is the standard error of 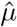, the approximate posterior distribution for OR is

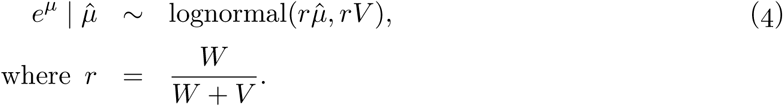

The logistic model includes the parameter of interest, *µ* = ln(OR), with the corresponding exposure value, *z*, as well as vectors of covariates and their respective coefficients, **x**^′^ and ***β***:

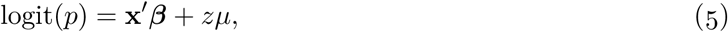

but the prior distribution is specified for *µ* only, thus the posterior distribution for *µ* is approximate. An estimator for ln(OR) can be obtained as an approximate posterior expectation of ln(OR), which is simply *r* 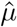.

The ABF method received much attention from the genetics community, in part due to simplicity of its calculation: the Bayes factor and the approximate posterior distribution for ln(OR) are computed based on the usual *Z*-statistic for ln(OR). The variance parameter *W* of a normal prior distribution for ln(OR) in ABF reflects the spread of the ln(OR) across SNPs in the genome.

### Simulation Study Setup

Given a population value of ln(OR) for a given SNP drawn from a prior distribution, one also needs to draw two other random population values, the probability of disease given a risk allele, Pr(*D*|*A*), and the allele frequency, Pr(*A*). Under Hardy-Weinberg equilibrium, Pr(*D*|*A*) is defined as an average over the risks for the genotypes that contain the allele *A*, i.e., *AA* and *AĀ*, and when the second allele in a genotype is also *A* with probability Pr(*A*), and *Ā* otherwise. The simulation setup can be summarized as follows:

1. For each SNP, draw ln(OR) from the prior distribution.
2. Draw population values of Pr(*D* | *A*) ∼ Beta(0.9, 2) and Pr(*A*) ∼ Uniform(0.1, 0.9).
3. Other population values can now be computed as follows:

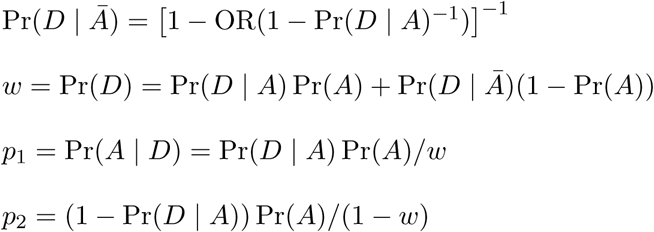
4. Data generating step. Draw two binomial samples of alleles for case and control individuals,

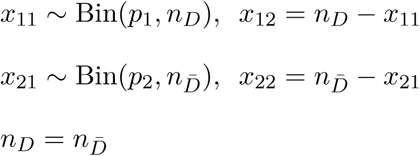 The total sample size in this model is the number of alleles (twice the number of individuals). Under the assumed Hardy-Weinberg equilibrium, analysis of allele counts approximates analysis of genotype counts by the usual trend test.^29^ Next, add 0.5 to the four counts, *n*_*ij*_ = *x*_*ij*_ + 0.5.

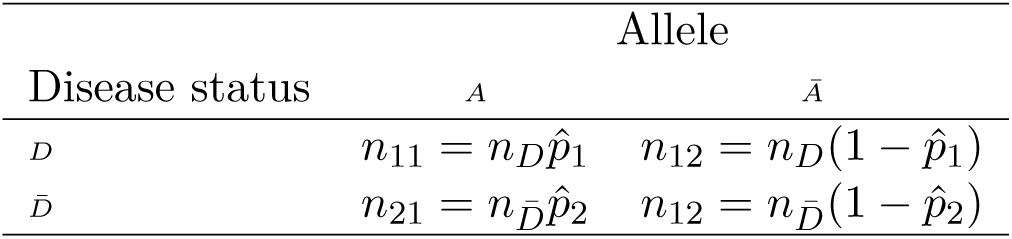 Based on the counts, compute *Z*-statistic as:

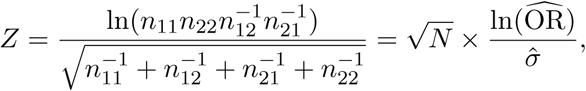

where

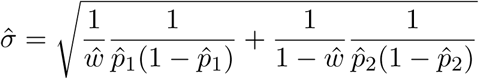 Sample proportion of cases, *ŵ*, is 1/2 by design, due to 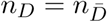.
5. Use Equ.2, to obtain the posterior probabilities and Equ. 3 to obtain the posterior expectation *δ*. The posterior estimate for ln(OR) is given by

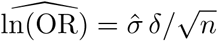

